# chromPlot: visualization of genomic data in chromosomal context

**DOI:** 10.1101/034108

**Authors:** Karen Y. Oróstica, Ricardo A. Verdugo

## Abstract

**Summary:** Visualizing genomic data in chromosomal context can help detecting errors in data generation or analysis and can suggest new hypotheses to be tested. Here we report a new tool for displaying large and diverse genomic data in idiograms of one or multiple chromosomes. The package is implemented in R so that visualization can be easily integrated with its numerous packages for processing genomic data. It supports simultaneous visualization of multiples tracks of data, each of potentially different nature. Large genomic regions such as QTLs or synteny tracts may be shown along histograms of number of genes, genetic variants, or any other type of genomic element. Tracks can also contain values for continuous or categorical variables and the user can choose among points, points connected by lines, line segments, barplots or histograms for representing data. chromPlot reads data from tables in BED format which are imported in R using its builtin functions. The information necessary to draw chromosomes for mouse and human is included with the package. Chromosomes for other organisms are downloaded automatically from the Ensembl website or can be provided by the user. We present common use cases here, and a full tutorial is included as the packages's vignette.

**Availability:** chromPlot is distributed under a GLP2 licence at Genomed Lab: http://genomed.med.uchile.cl.

## 1 INTRODUCTION

When analyzing large amounts of genomic data, visualization in chromosomal context can reveal errors in the data generation process or in a previous analysis step. It may also show chromosome-specific or genome-wide patterns that can suggest new hypotheses and the appropriate tests. We distinguish between two types of genomic visualization: local and global. Local visualization displays data in a region of one chromosome, with a resolution that may reach the basepair level. Global visualization displays data in whole chromosomes. There are powerful tools available for local visualization such GenomeGraphs (Durinck *et al*., 2009) and GenoPlotR (Guy *et al*., 2010) in R and the Integrative Genomics Viewer (IGV) (Thorvaldsdttir *et al*., 2013) written in Java. These software have become popular because of their flexibility to display any type of genomic data using customized plots. An equally flexible tool is needed for global visualization when analyzing large amounts of genome-wide data.

Global Visualization has been incorporated in websites of cross-species repositories such as Genome Graphs (Bioinformatics, 2015), or the Genome Decoration Page (NCBI, 2015). They have also been implemented in species-specific websites, such as or the Chromosome Visualization Tool (Cvit) for leguminous plants (Cannon & Cannon, 2011) and Gviewer for human, mouse, and rat (Twigger, 2009). There are also standalone tools that were developed with a specific purpose in mind. For instance, PhenoGram (Wolfe *et al*., 2013) is a program to represent genotype-by-phenotype associations. Quantsmooth creates a plotting area with a horizontal chromosome so that data can be added using genomic coordinates. It displays discrete and continuous data in the local view, and large segments in the global view. It cannot, however, bin data to represents large datasets as histograms in the global view. The IdeoViz R-package implements global views of continuous values as lines or bars for binned data (S & J, 2014). However, it cannot show large segments or individual points data, and it cannot plot on the chromosomal body, restricting the types of plots that can be created.

We introduce the chromPlot R-package, a generic tool for global visualization of genomic data. It can be used for any organism, even without a reference genome. It can represent heterogeneous tracts of information simultaneously; both large segments and small and numerous (millions) genomic elements. Data can be shown along either side of chromosomes or on the chromosomal body using a range of symbols.

## 2 SOFTWARE

chromPlot is fully coded in R and uses builtin function for data processing and plotting. It doesn’t have any other dependency than R itself. However, if GenomeGraphs is available, it can call it internally when the user wants to switch between global to local visualization. The package includes the data needed to plot ideograms for human and mouse. In addition, if the user wants to plot chromosomes from a different organism, tables can be downloaded from the UCSC genome browser and directly imported into chromPlot. Detailed instructions are provided in the package’s vignette.

The software is provided as and R-package, available from our website http://genomed.med.uchile.cl/software and has been submitted to Bioconductor.org for wide distribution.

## 3 PROGRAM OVERVIEW

### 3.1 Input data

chromPlot takes data as R objects of data.frame class. Tables must follow the BED format, which is commonly used to to share genomic annotations. The columns Chrom, Start, End are mandatory and column names must match these words exactly. A Name, Group, and Statistic column can be added to provide a genomic element ID, category, or continuous values when appropriate.

### 3.2 Usage

The user interacts with a single main function called `chromPlot`. Tracks of data are provided as named arguments. Argument names are specific for each type of data.

*Idiograms* chromPlot can show coloured bands on the chromosomal bodies. This can be used to create standard idiograms or pro display any other type of data assigned to multiple categories. Data is provided as a table to the band argument.

*Small elements* Four arguments are available for tracks of small genomic elements: annot, arraytxs, filtxs, seltxs. They are appropriate for elements like genes, array probes, small genetic variants, etc. By default, they are plotted as histograms when they are smaller than the size of bins (1Mb by default) but this can be changed by setting the noHist argument to TRUE. For drawing genes, it is possible to obtain them directly from the Ensembl repository. The user must provide an organism name as the org argument and annotBiomaRt=TRUE and the biomaRt package must be installed (Steffen Durinck & Huber, 2005). Otherwise, they must be provided as a table in the annot argument, but only one of these options can be used at a time.

*stackedBarplot* When genomic elements can be assigned to multiple categories, the used may display their frequency per bin as stacked barplots. The table with data provided to the stackedBarplot argument must contain a Group column.

*Large segments* the segment and segment2 arguments accept tracks of large segments. They are plotted as coloured lines on either side of the chromosomes. Optionally, the use may include up to two categories for each segment by providing the Group and Group2 columns with categorical values. Segments are be distinguished by color and a shape in the middle, which are labeled in a legend.

*Points and connected lines* The stats and stats2 arguments are used to display continuous data. Both tables must contain a Statistic column with continuous values to be plotted. By setting the argument statsTyp to ”I” or ”p”, the user can choose between connected lines or points.

## 4 RESULTS

chromPlot creates karyotype-like graphs and the data can be plotted on three areas along the x axis, to the left, to the right and on the chromosome. Multiple tracks may be overlaid or shown on different chromosomal sides. If multiple chromosomes are plotted, the canvas is divided into independent plotting areas. However, the same system of coordinates is used among chromosomes. Fifty four arguments are available for the user to generate a virtually infinite range of possible graphs. Here we list a short list of common use cases of the package.

*Synteny plots* To represent synteny between two organisms, the user’s data can be provided in AXT or BED format. The genomic regions of homology are represented by colored segments within chromosomes.

*Results from differential expression experiments* Histograms are useful to explores results from tests of differential expression in chromosomal context. This may reveal clusters of differential expression that may suggest effects from regional context such as chromatin state, epigenetics regulation, etc.

*QTL mapping* Genetic mapping detects association between genetic variation and phenotypes that can be represented as chromosomal segments. It is now possible to integrate such information with results from *omics* experiments, such as differential expression (Fig. 1).

**Fig. 1.**
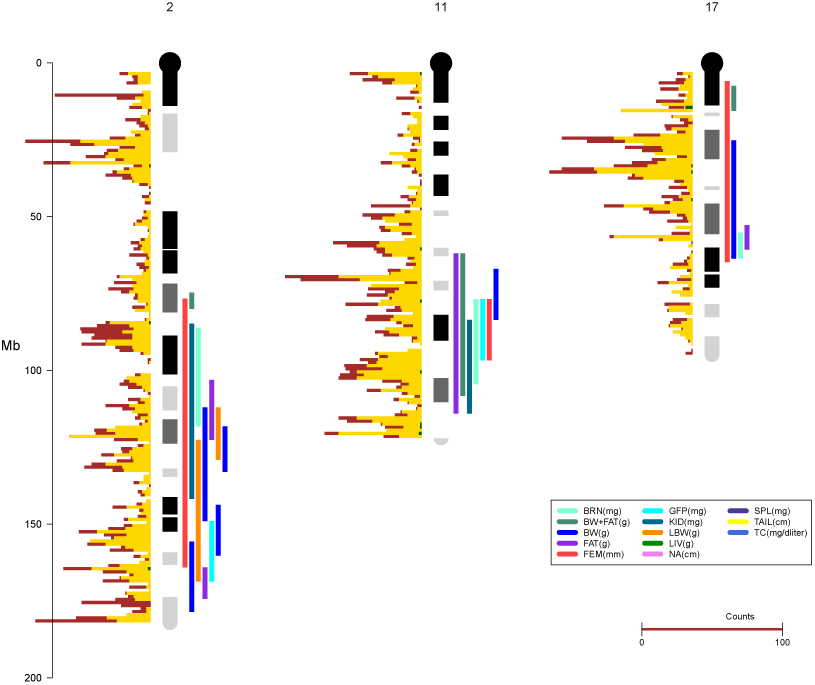
Genes selected by differential expression and identification of QTLs on chromosomes 2, 11 and 17. G banding is shown o chromosomal bodies. The color bars to the right of the chromosome are QTL. The number of genes in 1 Mbp bins along the chromosome is plotted as histograms to the left of each chromosome: brown, total number of genes; yellow, number of genes represented in the microarray; green, number of differentially expressed genes. The scale labeled Gene counts represents the number of genes. Details of the data in (Verdugo *et al*., 2010).

*GWAS* Genomewide association in outbred populations provide pointwise association between markers and phenotypes. These may be summarized as, for instance, average −log10(p) values in chromosomal bins. Similar plots may be created to summarize genomewide genetic divergence (e.g. Fst) or measures of positive selection. These applications are examples of point and connected-line plots.

*Genome sequencing* chromPlot can be used to quality control or summarizing results from whole-genome sequencing experiments. For instance, it is possible to plot genome coverage as connected-lines or number of detected variants as histograms.

## 5 DISCUSSION AND CONCLUSION

chromPlot is a generic and highly flexible tool for global visualization of genomic data. It’s able to plot large datasets for different scientific needs. Its been developed in R to facilitate inclusion in genomic workflows of data analysis. This feature makes chromPlot a more versatile software compared to other global visualization tools currently available.

## ACKNOWLEDGEMENT

We thank to Alejandro Blanco and Paloma Contreras for testing the package and providing useful comments that helped to improve the software.

*Funding:FONDECYT Initiation Project 11121666*.

## REFERENCES

Bioinformatics, UCSC Genome. 2015. Genome Graphs UCSC Genome Bioinformatics.https://genome.ucsc.edu/cgi-bin/hgGenome [Accessed: 2015-12-07].

Cannon, Ethalinda K. S., & Cannon, Steven B. 2011. Chromosome Visualization Tool: A Whole Genome Viewer. International Journal of Plant Genomics, 2011.

Durinck, Steffen, Bullard, James, Spellman, Paul T, & Dudoit, Sandrine. 2009. GenomeGraphs: integrated genomic data visualization with R. BMC Bioinformatics, 10(Jan.), 2.

Guy, Lionel, Roat Kultima, Jens, & Andersson, Siv G. E. 2010. genoPlotR: comparative gene and genome visualization in R. Bioinformatics, 26(18), 2334–2335.

NCBI. 2015. Genome Decoration Page. http://www.ncbi.nlm.nih.gov/genome/tools/gdp/ [Accessed: 2015-12-07].

S, Pai, & J, Ren. 2014. IdeoViz: Plots data (continuous/discrete) along chromosomal ideogram. R package version 1.4.0.

Steffen Durinck, Yves Moreau, Arek Kasprzyk Sean Davis Bart De Moor Alvis Brazma, & Huber, Wolfgang. 2005. BioMart and Bioconductor: a powerful link between biological databases and microarray data analysis. Bioinformatics, 21(16), 3439–3440.

Thorvaldsdttir, Helga, Robinson, James T., & Mesirov, Jill P. 2013. Integrative Genomics Viewer (IGV): high-performance genomics data visualization and exploration. Briefings in Bioinformatics, 14(2), 178–192.

Twigger, Simon. 2009. Flash GViewer. http://gmod.org/wiki/Flash_GViewer [Accessed: 2015-12-07].

Verdugo, Ricardo A., Farber, Charles R., Warden, Craig H., & Medrano, Juan F. 2010. Serious limitations of the QTL/Microarray approach for QTL gene discovery. BMC Biology, 8(1), 96.

Wolfe, Daniel, Dudek, Scott, Ritchie, Marylyn D, & Pendergrass, Sarah A. 2013. Visualizing genomic information across chromosomes with PhenoGram. BioData Mining, 6(Oct.), 18.

